# Quantitative imaging of three-dimensional fiber orientation in the human brain via two illumination angles using polarization-sensitive optical coherence tomography

**DOI:** 10.1101/2023.10.20.563298

**Authors:** Chao J. Liu, William Ammon, Robert J. Jones, Jackson C. Nolan, Dayang Gong, Chiara Maffei, Brian L. Edlow, Jean C. Augustinack, Caroline Magnain, Anastasia Yendiki, Martin Villiger, Bruce Fischl, Hui Wang

## Abstract

The accurate measurement of three-dimensional (3D) fiber orientation in the brain is crucial for reconstructing fiber pathways and studying their involvement in neurological diseases. Optical imaging methods such as polarization-sensitive optical coherence tomography (PS-OCT) provide important tools to directly quantify fiber orientation at micrometer resolution. However, brain imaging based on the optic axis by PS-OCT so far has been limited to two-dimensional in-plane orientation, preventing the comprehensive study of connectivity in 3D. In this work, we present a novel method to obtain the 3D fiber orientation in full angular space with only two illumination angles. We measure the optic axis orientation and the apparent birefringence by PS-OCT from a normal and a 15 deg tilted illumination, and then apply a computational method yielding the 3D optic axis orientation and true birefringence. We verify that our method accurately recovers a large range of through-plane orientations from -85 deg to 85 deg with a high angular precision. We further present 3D fiber orientation maps of entire coronal sections of human cerebrum and brainstem with 10 μm in-plane resolution, revealing unprecedented details of fiber configurations. We envision that further development of our method will open a promising avenue towards large-scale 3D fiber axis mapping in the human brain and other complex fibrous tissues at microscopic level.

## 1. Introduction

The network of fibers that constitute the white matter in the human brain provides a complex set of connections underlying sophisticated functions. Deciphering the structural connectivity of the white matter is thus crucial to understand human brain function and behavior. Diffusion weighted magnetic resonance imaging (dMRI) allows fiber pathways to be imaged *in vivo* and has advanced significantly in recent years due to initiatives such as the Human Connectome Project ^1,2^. However, because of the limited resolution of MRI techniques, microscopic methods are still needed to obtain the ground truth of fiber architecture, particularly in the regions where multiple fiber bundles intersect ^3^. Optical methods have used various contrast mechanisms to image myelinated axons at micrometer resolution, such as fluorescent labeling ^4^, third harmonic generation ^5^ and anti-stokes Raman scattering ^6^. However, these techniques still require the fiber orientation to be estimated from the acquired images, using techniques such as structure tensor analysis, which has been applied to histological staining ^7,8^, fluorescence microscopy ^9^, and label-free optical coherence tomography (OCT) images ^10^.

Imaging techniques using light polarization take the advantage of the birefringent nature of the myelin sheath surrounding axons to generate quantitative and label-free contrasts for myelinated fibers ^11^. The fiber geometry causes optical anisotropy with negative birefringence and an optic axis orientation aligned with the physical fiber direction. In recent years, prominent methods such as polarized light imaging (PLI) ^12^ and automatic serial sectioning polarization-sensitive OCT (PS-OCT) ^13,14^ have been applied to map microscopic fiber orientation in large-scale brain samples. PLI uses multiple oblique views to derive the fiber orientations in human brain sections, in which physical cutting of the sample into 50-100 μm thick slices is a prerequisite ^15^. In serial sectioning PS-OCT, the blockface of the sample is imaged prior to sectioning, thus avoiding section-to-section distortions. The conventional optic axis orientation measurement of PS-OCT from one illumination angle does not describe the 3D fiber orientation, but rather its projection onto a plane orthogonal to the beam direction. One approach to extract the through-plane orientation is to use multiple illumination angles. The in-plane optic axis orientations obtained by different illumination angles allows for the calculation of the through-plane orientation via a geometrical derivation. Our previous study suggested a minimum of three beams to accurately recover the 3D orientation of muscle fibers ^16^. Similarly, various illumination configurations have been proposed in PS-OCT and validated in birefringent samples of tendons and muscles ^16–19^, using widely spaced imaging directions, and either excluding or assuming optic axes co-planar to the illumination beam directions. Furthermore, 3D fiber orientation measurements using PS-OCT in the human brain have not been reported.

In this work, we present a novel computational method to estimate 3D axis orientation in human brain tissue via only two illumination beams separated by a small angle of 15 deg. This approach formulates a birefringence vector composed of optic axis orientation and true birefringence, which is estimated to match to the two measured birefringence values and in-plane optic axis orientations. We first validate the performance of our approach with a uniform section of a corpus callosum sample. We then illustrate examples of the 3D fiber orientation in large human brain samples, including coronal sections of a whole hemisphere and a medulla sample.

## 2. Methods

## 2.1 3D orientation reconstruction algorithm

Previous PS-OCT work has reported that the true 3D optic axis can be recovered by the measurements of apparent birefringence ^17^ or optic axis orientations ^16^ at variable illumination angles. Both metrics show a dependence on the illumination angles and the measurements from different incident angles are linked by the through-plane orientation of the fibers. Geometrically, the 3D optic axis lies along the intersection of multiple planes, each of which is formed by an incident beam and its corresponding optic axis orientation [20]. However, it is necessary to also account for the effective birefringence values, in order to manage the case when the optic axis is co-planar to the illumination beams, and hence the two planes coincide. We utilized these physical relationships to develop our algorithm.

To estimate the 3D optic axis via two illuminations, we define a birefringence vector Δ***n*** = Δ*n*[*o_x_ o_y_ o_y_*]^*T*^, where the magnitude Δ*n* is the true birefringence value and the unit vector [*o*_*x*_ *o*_“_ *o*_#_]^*T*^ represents the 3D orientation. The in-plane (Θ) and through-plane (*α*) orientations are obtained by vector calculus of [*o*_*x*_ *o*_“_ *o*_#_]^*T*^ in spherical coordinates. Let ***p*** be the unit vector parallel to the illumination beam. We use two illumination beams, one along the z-axis, ***p***_2_ = [0 0 1]^*T*^, and one rotated about the y-axis by Ω, ***p***_2_ = [*sin*Ω 0 *cosΩ*]^*T*^. We obtain two measurements of the apparent optic axis orientation Θ_’_ and Θ_(_, and two measurements of the apparent birefringence Δ*n*′_’_ and Δ*n*′_(_. This yields an apparent birefringence vector Δ***n***′_2_ = Δ*n*′_’_[*cos n*Θ_’_ 0] ^*T*^ under the normal illumination ***p***_2_ (Fig. 1, left panel) and Δ***n***′_2_ = Δ*n*′_(_[*cosΩcos*Θ_2_ *sin*Θ_2_ -*sin*Ω*cos*Θ_2_]^*T*^ under the tilted illumination ***p***_2_ (Fig. 1, right panel). It is noted that the apparent birefringence values in each of the two frames are related to the true birefringence ^20^ as Δ*n*′_’_ = Δ*n sin*^(^Ψ_’_ and Δ*n*′_(_ = Δ*sin*^(^Ψ_(_, where Ψ_’_ and Ψ_(_ are the polar angles (*π*⁄2 − *α*) in the two illumination frames.

**Fig. 1.**
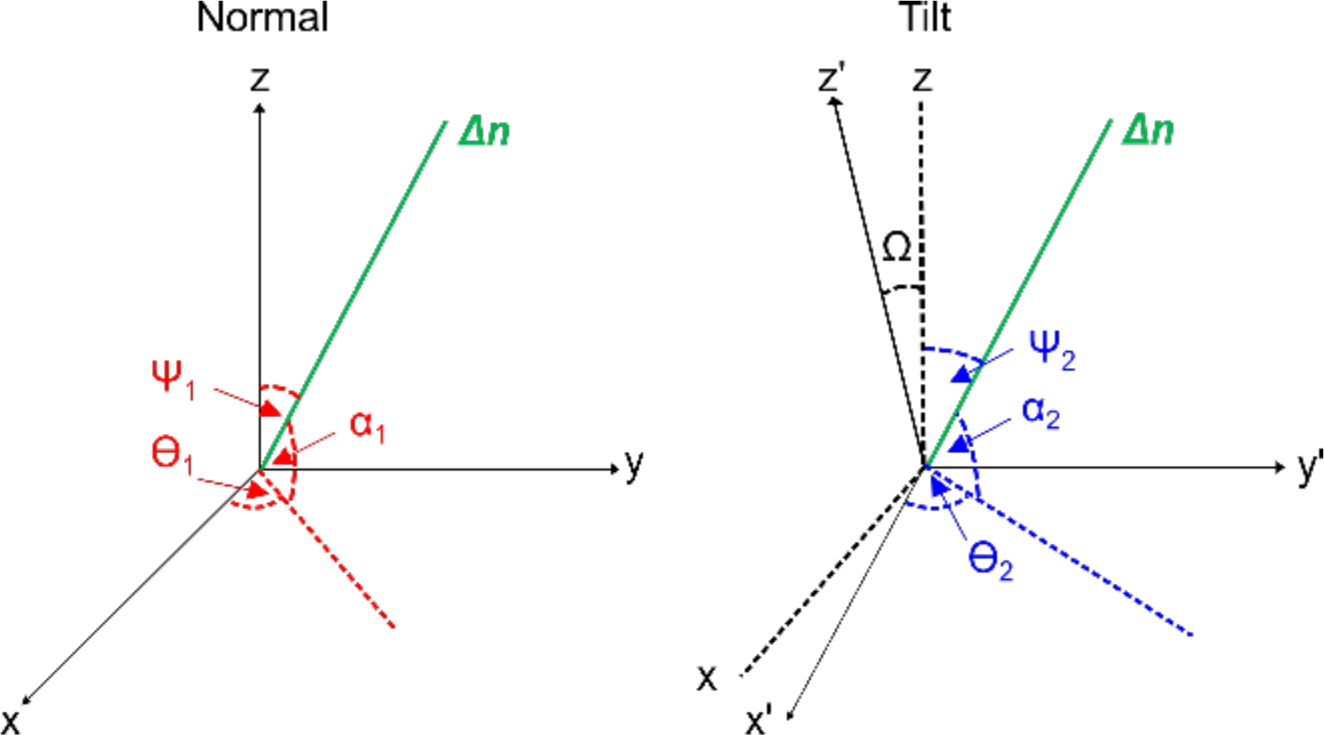
The coordinate system and angle definition in the normal (left) and tilted (right) illumination incidences.

We now define an estimated birefringence vector 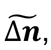 which in turn defines its estimated in-plane orientations and polar angles, 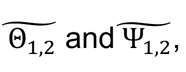 respectively, in the two frames. Therefore, the estimated apparent birefringence vector 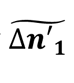 under the normal illumination ***p***_1_ can be written as,

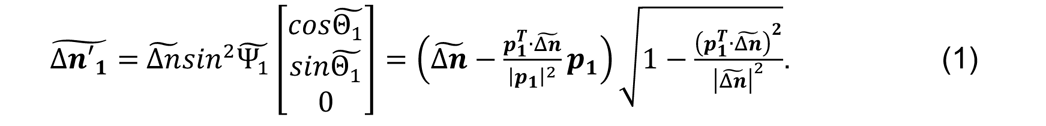

where the expression in the parenthesis projects the birefringence vector into the plane orthogonal to 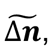 thereby scaling it by 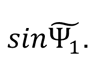 The square root adds the required second 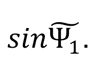 Similarly, the estimated apparent birefringence vector 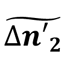 under the tilted illumination ***p***_2_ can be written as,

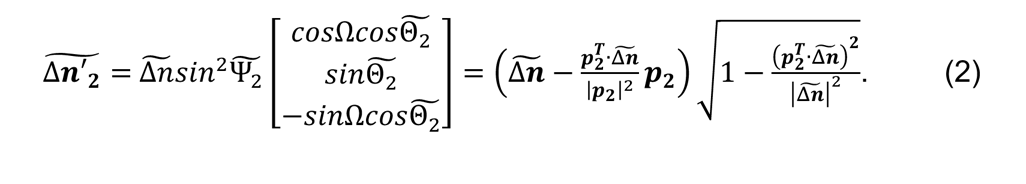

We then formulate an optimization problem in Eq. 3 that finds the optimal birefringence vector 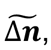 such that the sum of the mean square errors between the estimated and the measured apparent birefringence vectors under the two illuminations is minimum:

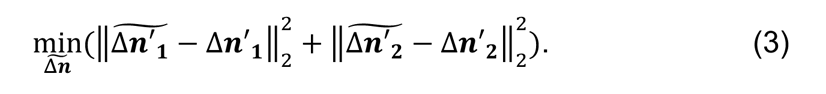

The above computational model works well for the majority of the fiber orientations. However, a problem arises when applying Eqs. 1 – 3 to the fibers with in-plane orientations running along the y-axis (i.e., Θ_’_ ≈ Θ_(_ ≈ −90/90 *deg*). Since the optic axis measurement ranges between -90 deg and 90 deg, the distribution of the noise at Θ ≈ −90/90 *deg* forms a U-shape (bounded by -90 deg and 90 deg, see details in Fig. S1), which deviates from the close to normal distribution of the noise at intermediate angles in Eq. 3. Consequently, the optimization output is sub-optimal. To overcome this problem, we interchanged the x-axis and y-axis for those fibers in post-processing, to effectively convert the optic axis orientation to close to 0 deg. As a result, the noise distribution is returned back to an approximately normal one. The tilted illumination in the swapped xy-axis is equivalent to tilting about x-axis by Ω, which yields the apparent birefringence vector as Δ***n***′_2_ = Δ*n*′_(_[cos Θ(*o*cosΩ_2_sinΘ(*sinΩsinΘ*_2_] ^*T*^. Correspondingly, the estimated apparent birefringence vector is written as,

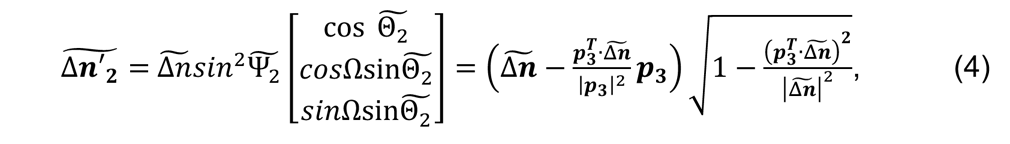

where ***p*** = [0 − *n*Ω *o*Ω]^*T*^ is the new direction vector of the tilted illumination beam about the x-axis. We found the standard deviation of the orientation along one A-line was 26.7 deg (average of 100 A-lines). Therefore, we combine the two sets of equations and adapt a criterion that Eqs. 1, 2, and 3 are used to retrieve fibers with optic axis |Θ_’_| < 45 and Eqs. 1, 3, and 4 are used to estimate fibers with optic axis |Θ_’_| ≥ 45 *deg* (see section 3.2).

### 2.2 Human brain samples

We used samples excised from two neurologically normal brains from the Massachusetts General Hospital (MGH) Autopsy Suite for this study (age at death: 70 years and 46 years; 1 male and 1 female) and one neurologically normal brainstem sample from a subject enrolled in the Université de Tours body donation program (age at death: 76 years, male). Before death, the participants provided written proof of their willingness to become body donors and consented to the use of their bodies for educational or research purposes. The post-mortem intervals did not exceed 24 hours. The brains were fixed by immersion in 10% formalin and were cut into smaller blocks. We cut two blocks in the corpus callosum region from one of the MGH samples and one coronal slab from the other MGH sample. We applied refractive index matching to all the samples by incubations in 2,2-thiodiethanol solutions as described in our previous work ^21^. The *ex vivo* imaging procedures are approved by the Institutional Review Board of the MGH.

### 2.3 System, data acquisition and analysis

We used a custom-built automatic serial sectioning PS-OCT (as-PSOCT) system ^13,21^ to validate our 3D orientation method. The system consists of a spectral domain PS-OCT centered at 1300 nm, motorized xyz translational stages, and a vibratome for tissue sectioning. The PS-OCT is a free-space system. A quarter wave plate is placed in the sample arm to ensure circularly polarized light on the sample. Using a Jones matrix formalism, the retardance and the optic axis orientation are obtained by the amplitude and the phase of the complex signals, respectively, on the two polarization channels. Automatic imaging and sectioning of brain blocks is controlled by custom-built software for coordinating data acquisition, xyz stage translation, and vibratome sectioning. The samples were first flat faced with the vibratome and then imaged using a scan lens (OCT-LSM3, Thorlabs, Newton, NJ). The range of imaging depth was 2.6 mm with an axial resolution of 4.2 μm in tissue.

We imaged the sample with normal illumination and illumination tilted by 15 deg about the y-axis. To implement the tilted illumination, we either mounted the sample on a rotation stage or incorporated a rotation stage into the PS-OCT scan head. The physical effect of tilting the beam or the sample is the same ^16^. For the normal illumination, one volumetric acquisition covered a field of view (FOV) of 3.5 mm × 3.5 mm. The FOV was 3.5 mm × 2 mm for the 15 deg tilted illumination. Automatic tile-scan was used to cover the whole sample, with a 20% overlap between tiles in each direction of the raster scan. For volumetric imaging of large-scale samples, a 150 µm thick slice was removed from the tissue surface by the vibratome to expose the deeper region until the whole volume was imaged.

We obtained measurements of the apparent birefringence and the optic axis orientation from the two illumination angles. The apparent birefringence was calculated by performing a linear regression of the retardance profile along 150 μm in depth and extracting the slope of the linear fit. *En-face* optic axis orientation images were determined by the peak of a histogram formed by binning the measured orientation values into 5 deg intervals along 150 μm depth. The apparent birefringence and the *en-face* orientation data were then used to compute 3D axis orientation and true birefringence (see section 2.1).

### 2.4 Image registration between the normal and titled illuminations

We applied a demon registration algorithm ^22,23^ to register the images between the normal and titled illumination incidences. We first registered the apparent birefringence maps from the two incidences and obtained a transformation field. Then we applied this transformation field to register the optic axis orientation maps.

### 2.5 Estimation of 3D orientation

The optimization algorithm takes the registered *en-face* orientation (Θ_’_ and Θ_(_) and apparent birefringence (Δ*n*′_’_ and Δ*n*′_(_) from normal and tilted incidences as inputs, converts them into the birefringence vector Δ***n***′_2_ and Δ***n***′_2_, and outputs the estimated Δ**n**, which yields the true birefringence value Δ*n* and the 3D orientation vector [*o*_*x*_ *o*_“_ *o*_#_]^*T*^ (see details in Section 2.1). The 3D orientation vector is further converted into in-plane and through-plane orientations for better visualization. We used the function *fminsearch* in Matlab (Mathworks, Natick, MA) to minimize the difference between the estimated and the measured birefringence vectors. The initialization values were set to be [0, 0, 0] and the optimization stopped based on an error tolerance of 1e-20 in the *fminsearch* function.

### 2.6 Quantifying fiber orientation distribution

To quantitatively analyze the orientations obtained from the uniform corpus callosum sample, we constructed fiber orientation distributions (FODs) by generating histograms of the orientation map in every 0.5 mm x 0.5 mm region using a bin width of 5 deg. We found the primary peak and full width at half maximum (FWHM) of each FOD, then calculated the averaged peak direction and FWHM over the FODs in a 2.5 mm × 2.5 mm or 3 mm × 3 mm region of interest (ROI). Pixels with low birefringence values were excluded from the analysis.

### 2.7 Diffusion weighted MRI

The sample of coronal slab was packed in fomblin and scanned in a small-bore 4.7 T Bruker BioSpin MRI scanner (maximum gradient strength 660 mT/m) using a diffusion-weighted 3D echo-planar imaging sequence with the following parameters: 0.25 mm isotropic resolution, TR = 500 ms, TE = 47.8 ms, 20 segments, 5 averages, δ = 15 ms, Δ = 19 ms, b-value = 4,000 s/mm^2^, one b=0 volume and 12 volumes with non-colinear gradient directions. After denoising ^24^ and correction for eddy-current distortions ^25^, deterministic tensor tractography ^26^ was performed, seeding in every white-matter voxel.

## 3. Results

### 3.1 Range and accuracy of through-plane axis orientation measurement

We used a section of a human corpus callosum in which the fibers run in a mostly uniform direction to validate the 3D fiber orientation estimation. To create different through-plane angles, we positioned the sample on horizontal surface (0 deg) and two triangular wedges made of agarose gel (-30 and 30 deg, Fig. 2a), and obtained PS-OCT measurements from the normal and 15 deg tilted illuminations. The long axis of the fibers was positioned along the x-axis of the laboratory frame. Our optimization yielded the same in-plane orientations (Fig. 2b), regardless of the sample inclination, whereas differentiated through-plane orientations (Fig. 2c). We found the mean of the through-plane FODs to be 9.2 deg, -26.9 deg and 37.7 deg, respectively, on 0 deg surface (no wedge), -30 deg wedge, and 30 deg wedge (Fig. 2d), successfully revealing the angular differences between the three inclinations. In particular, our method was able to distinguish the positive and negative through-plane orientations (green vs. magenta wedges in Fig. 2). The consistent in-plane orientations illustrates the reliability of our estimations.

**Fig. 2.**
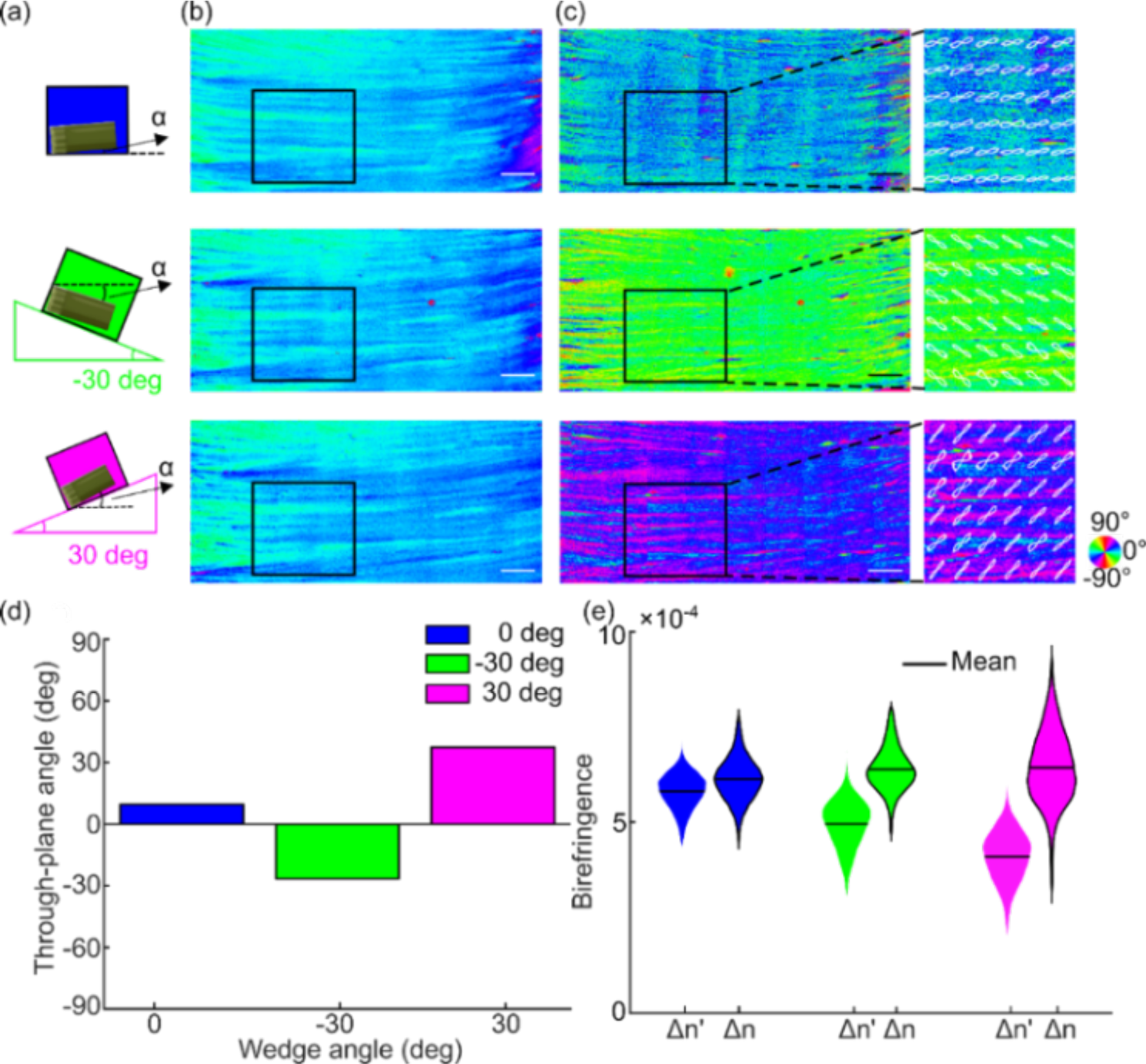
Characterization of through-plane orientation measurement for different inclination angles. (a) A cartoon illustrating the corpus callosum sample positioned on 0 deg (no wedge), -30 deg wedge, and 30 deg wedge to create different inclination angles. *α*: inclination angle. (b) The in-plane orientation images of the sample on three wedge scenarios. (c) The through-plane orientation images of the sample on three wedge scenarios. The enlarged boxes of the ROI show the FODs overlaid on the orientation image. The angles of orientation in (b) and (c) are indicated by the color wheel. (d) The average through-plane orientation of the FODs within the ROIs in (c). (e) The violin plots of average apparent (Δ*n*′) and true birefringence (Δ*n*) within the ROI in (c). The mean value of the birefringence is indicated by the black line. Scale bars: 1 mm.

We also investigated the estimation of true birefringence. We computed the mean of the apparent birefringence Δ*n*′ and true birefringence Δ*n* within the ROIs from the three inclination setups. The apparent birefringence (5.8 x 10^-4^, 4.9 x 10^-4^ and 4.1 x 10^-4^) reduced in the ±30 deg inclination setups, showing the dependency on the through-plane angle (Fig. 2e). In contrast, we successfully obtained the same true birefringence across the three setups regardless of the different through-plane angles (the mean value: 6.2 x 10^-4^), which captures the underlying myelin content in this corpus callosum sample.

We then used another block of the corpus callosum sample to characterize the range of our through-plane orientation measurement. We performed two independent experiments on different regions of the sample. The sample was positioned on a tilting stage to artificially vary the through-plane angle. In the first experiment, the stage was tilted counterclockwise from 0 deg to 50 deg in 10 deg steps. Our method successfully measured the angular differences between consecutive tilting angles (Fig. 3a). When the stage was positioned at 0 deg, the through-plane orientation of the fibers was measured to be 18.6 deg (averaged FODs, Fig. 3). As the tilting angle of the stage increased up to 50 deg, the mean FOD of the through-plane orientation was estimated to be 73.6 deg (Fig. 3b). A linear fitting of the estimated through-plane orientation with respect to the tilting angle revealed a slope of 0.97. In the second experiment, we imaged another region of the sample by tilting the stage at 10 deg, 30 deg and 50 deg angles and estimated the corresponding through-plane orientation. The slope of a linear fitting between the estimation and the pre-set tilting angle was 1.21. The two experiments characterized the angular range and precision of the through-plane orientation. It is noted that the through-plane orientation in the two experiments was different when there was no tilt in the stage, revealing the regional heterogeneity of the fiber tracts in corpus callosum. The results showed that we can accurately measure the through-plane orientation up to 85 deg. As Fig. 2 showed that we could differentiate the sign of the through-plane orientation, these ensemble of result suggested that our method is able to recover the full range of the through-plane fiber orientation in the human brain.

**Fig. 3.**
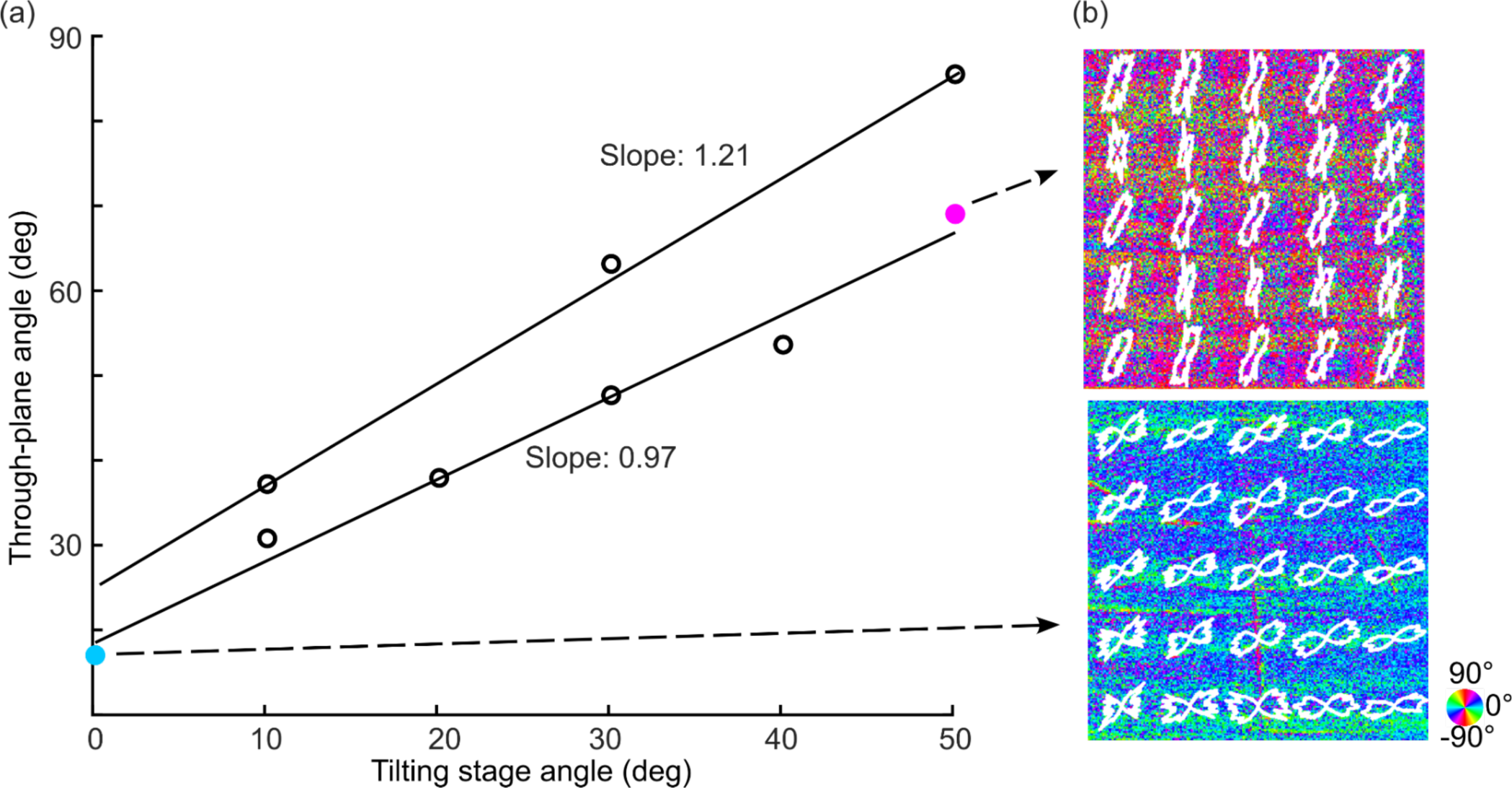
Characterization of the range of through-plane orientation estimation. (a) Two independent through-plane orientation measurements at different tilting stage angles. (b) Representative images of through-plane orientation when the tilting stage was set at 0 deg and 50 deg, with FODs overlaid. The angles of orientation are indicated by the color wheel. The sample size in (b) is 2.5mm x 2.5mm.

### 3.2 Accuracy of through-plane axis under variety of in-plane orientations

This section validates the estimation of through-plane axis under a variety of in-plane orientation scenarios. We rotated a corpus callosum sample counterclockwise from 0 to 90 deg in the XY plane by 30 deg steps and obtained measurements via the normal and 15 deg tilted illuminations (Fig. 4). Our method estimated the in-plane orientations accurately (Fig. 4a) when the sample was oriented at -90, -60, -30 and 0 deg. Meanwhile, the through-plane orientation estimate was consistent among the different in-plane orientation setups (Fig. 4b). To evaluate our method quantitatively, we analyzed the FODs of a 3 mm x 3 mm ROI as shown in Fig. 4a and Fig. 4b. The mean in-plane orientation estimation computed from the FOD was -82.8, - 57.2, -24.8 and 8.5 deg (Fig. 4c). Note that there was a small angular offset between the long axis of the fibers and the x-axis of the laboratory frame. The through-plane orientation estimation remained consistent at 2.8, -1.2, -2.4 and -1.6 deg across the different in-plane orientation settings (Fig. 4d). This suggests that our method can measure the 3D axis of all fibers accurately regardless of their orientation in the XY plane.

**Fig. 4.**
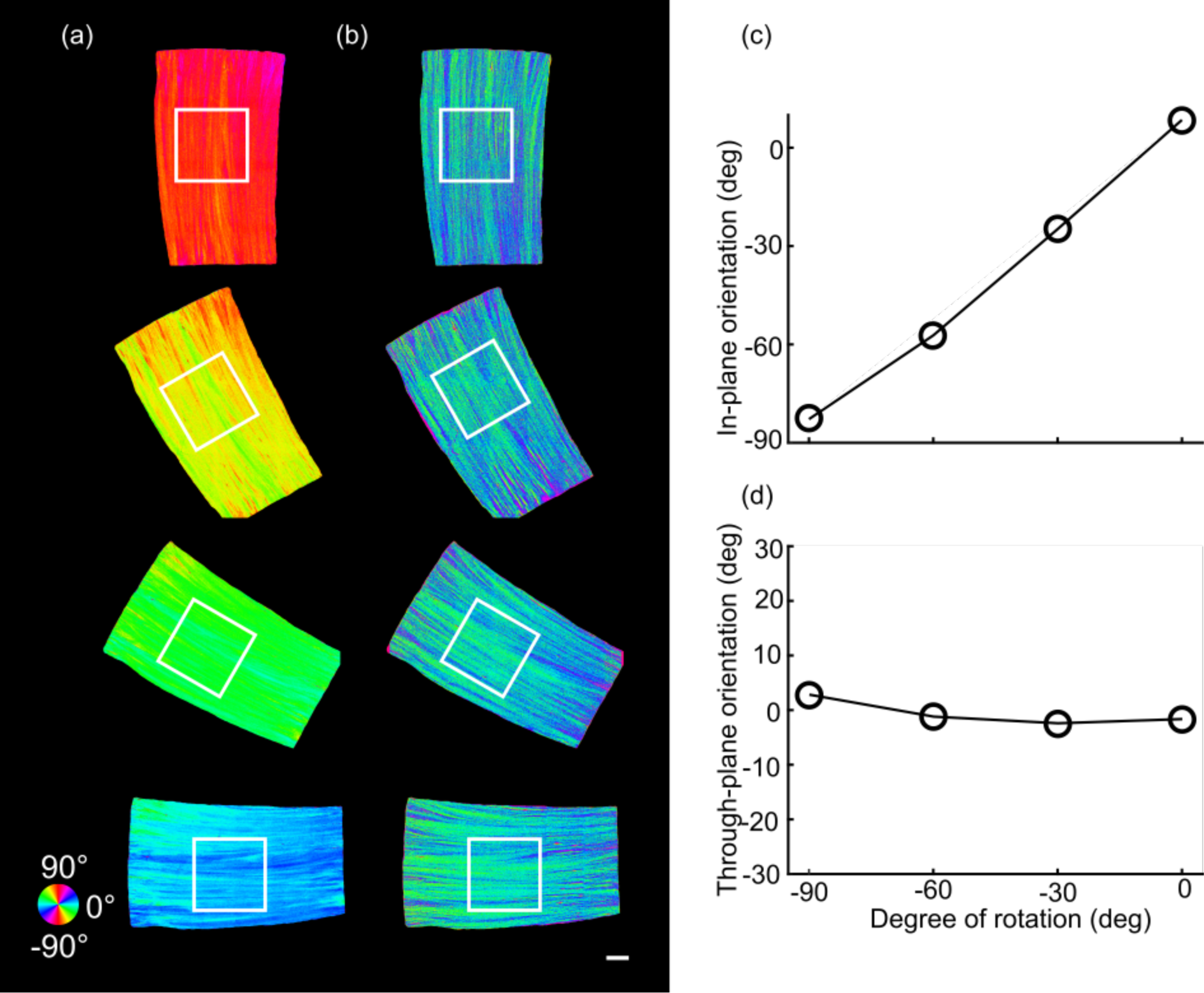
Characterization of through-plane orientation measurement at four different in-plane orientation angles on a corpus callosum sample. (a) The estimated in-plane orientation images at in-plane rotations of -90 deg, -60 deg, -30 deg, and 0 deg. (b) The corresponding through-plane orientation images. The black boxes indicate the ROIs to obtain the FODs. The angles of orientation in (a) and (b) are indicated by the color wheel. The average in-plane (c) and through-plane (d) orientation of the FODs within the ROIs shown in (a) and (b) with respect to the degree of rotation, separately. Scale bar: 1mm.

We demonstrated the advantage of the proposed method using an xy-axis swap criteria at 45 deg (Fig. 4, also see section 2.1), by comparing to the results without xy-swap (using Eq. 1, 2, 3 for all in-plane orientations). In the latter case, both the in-plane (Fig. S2a) and through-plane (Fig. S2b) orientation showed high errors at -90 deg rotation (see Fig. S1 for details). We also compared our optimization result against an earlier method using a geometric derivation (Fig. S2c, see details in reference ^16^). Using only one tilted beam about the y-axis, the geometric derivation method resulted in errors in through-plane estimation as the in-plane angle reached -60 deg. The error continued to grow as the in-plane angle increased to -90 deg.

## 3.3 3D fiber orientation map of a coronal section in the human brain

To investigate the feasibility of 3D orientation measurements in a large-scale sample, we imaged a coronal section of a human brain hemisphere, with a surface area of ∼6×9 cm^2^. We first acquired diffusional MRI data of the sample. For PS-OCT imaging, the tissue surface was flat-faced, and imaged with both normal and 15 deg tilted illuminations to recover the 3D orientations. We generated the in-plane and through-plane orientation maps from PS-OCT, and 3D orientation maps of the same plane both from PS-OCT and diffusion MRI (Fig. 5).

**Fig. 5.**
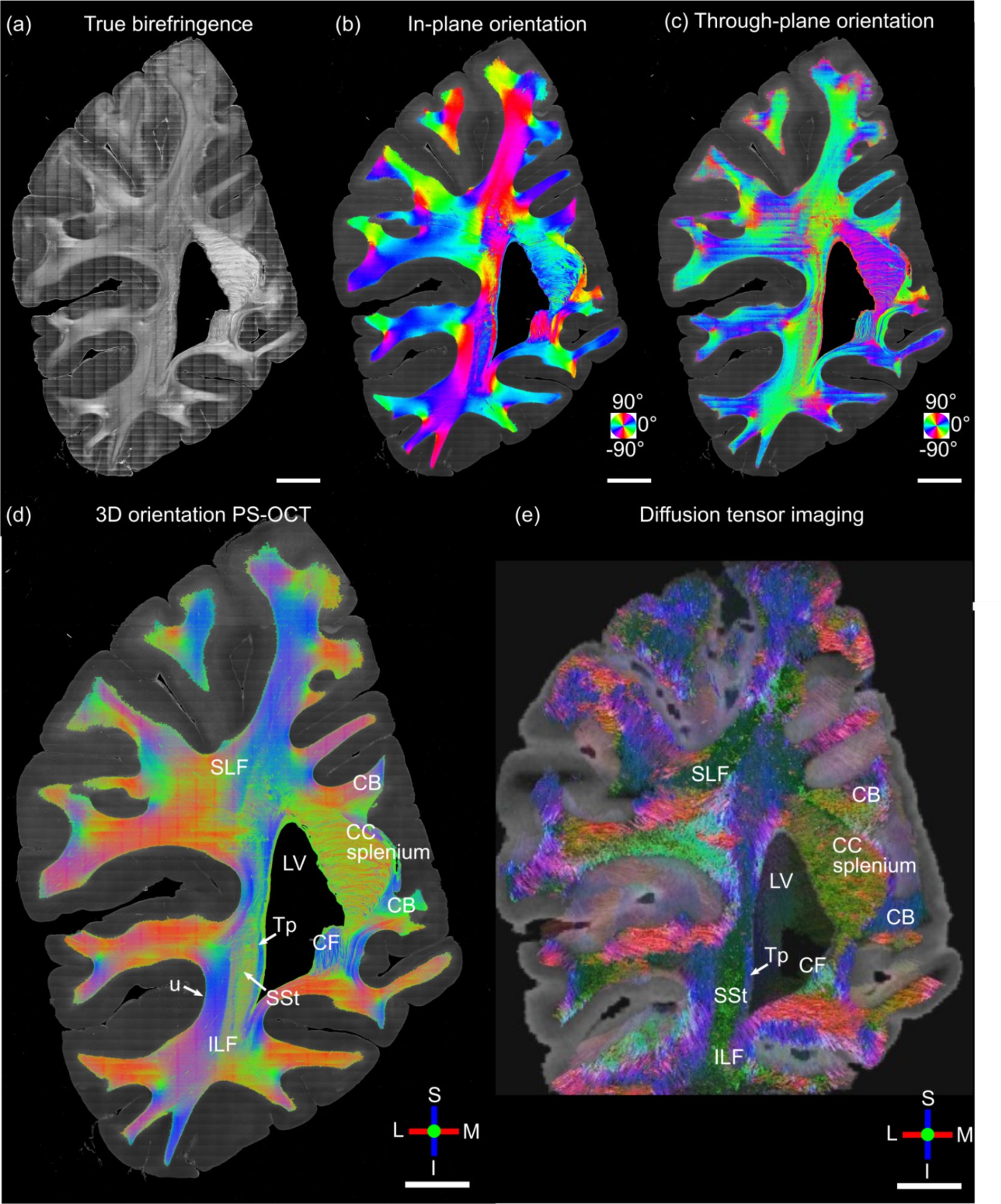
3D fiber orientation maps of the coronal slice sample. (a) True birefringence, (b)in-plane orientation, and (c) through-plane orientation. The in-plane and through-plane orientations are color-coded as indicated by the color wheels. 3D orientations are color-coded as indicated by the red (left/right), green (anterior/posterior) and blue (inferior/superior) orthogonal axes. Several anatomical structures are labeled in (d) and (e): LV, lateral ventricle; CC splenium, splenium of the corpus callosum; Tp, tapetum; SSt, sagittal stratum; ILF, inferior longitudinal fasciculus; u, ufibers. L: lateral; M: medial; S: superior; I: inferior. Scale bars: 10 mm.

Overall, the 3D orientation map of PS-OCT (Fig. 5d) is consistent with diffusion MRI (Fig. 5e) and the known neuroanatomy of white matter architecture in this region (Fig. 5) ^27^. In the 3D orientation map, medial-lateral/right-left (red) and inferior-superior (blue) directions are in-plane and anterior-posterior (green) is through-plane. The splenium of the corpus callosum is situated on the superiomedial border of the lateral ventricle and contains highly birefringent fibers oriented in the anterior-posterior (green) and medial-lateral (red) directions. Lateral to the lateral ventricle, three white matter tracts could be clearly delineated. The tapetum (Tp) borders the lateral edge of the lateral ventricle and contains fibers coursing inferior-superior (blue). The sagittal stratum (SSt) sits lateral to the Tp and contained fibers running primarily anterior-posterior (green). Lateral to the SSt, the inferior longitudinal fasciculus (ILF) consists of mostly inferior-superior oriented fibers (blue). Throughout the superficial white matter, short association fibers (i.e., U-fibers), are seen wrapping around the sulci and extending down the gyri into gray matter. Gyral fibers show predominantly in-plane orientations (red, blue), while small bundles of through plane fibers (green) are consistently observed at the base of sulci.

The high-resolution 3D fiber orientation map of PS-OCT depicts a diverse, heterogeneous organization of fiber bundles in deeper white matter. The smooth transition of the colored fiber orientation along tracts indicates realistic continuity that one might expect in the densely packed fascicular highways of the deep white matter.

## 3.4 3D fiber orientation map of a human brainstem

The micrometer resolution PS-OCT reveals complex configurations of small fiber tracts in brain regions beyond the ability of diffusion MRI. Particularly in the brainstem, compact fiber tracts are interconnected among small nuclei in all directions. To demonstrate the advantages of 3D axis orientation by PS-OCT, we imaged a slab of a human brainstem at the level of the medulla oblongata. The tissue surface was flat-faced, and imaged with both normal and 15 deg tilted illumination as described previously. The brainstem was positioned in an axial plane, i.e., left-right (red), and anterior-posterior (green) being the x and y axis of our laboratory frame, respectively (Fig. 6). It is well known that there are large numbers of fibers running through the axial plane in the brainstem. We annotated some of the major structures on the high resolution images based on the Paxinos Atlas (Fig. 6e) ^28^ and inspected the orientation of the fiber tracts. For example, the direction of internal arcuate fibers seen in the reticular formation (RF) running from the cuneate (Cu) and gracile nuclei (not shown in Fig. 6) to the contralateral medial lemniscus (ml), is consistent with the mostly in plane fibers in the RF and ml. The amiculum of the olive (ami) is a capsule of myelinated fibers surrounding the inferior olivary (IO) running both in-plane and through-plane. The spinal trigeminal tract (sp5), as well as the inferior cerebellar peduncle (icp), run inferior to superior, connected with other structures via numerous input and output fibers throughout their length in the axial plane. Finally, two other main fiber tracts, the pyramidal (pyr) and solitary tract (sol) run inferior to superior and thus appear mostly blue in the color rendering of the 3D fiber orientation.

**Fig. 6.**
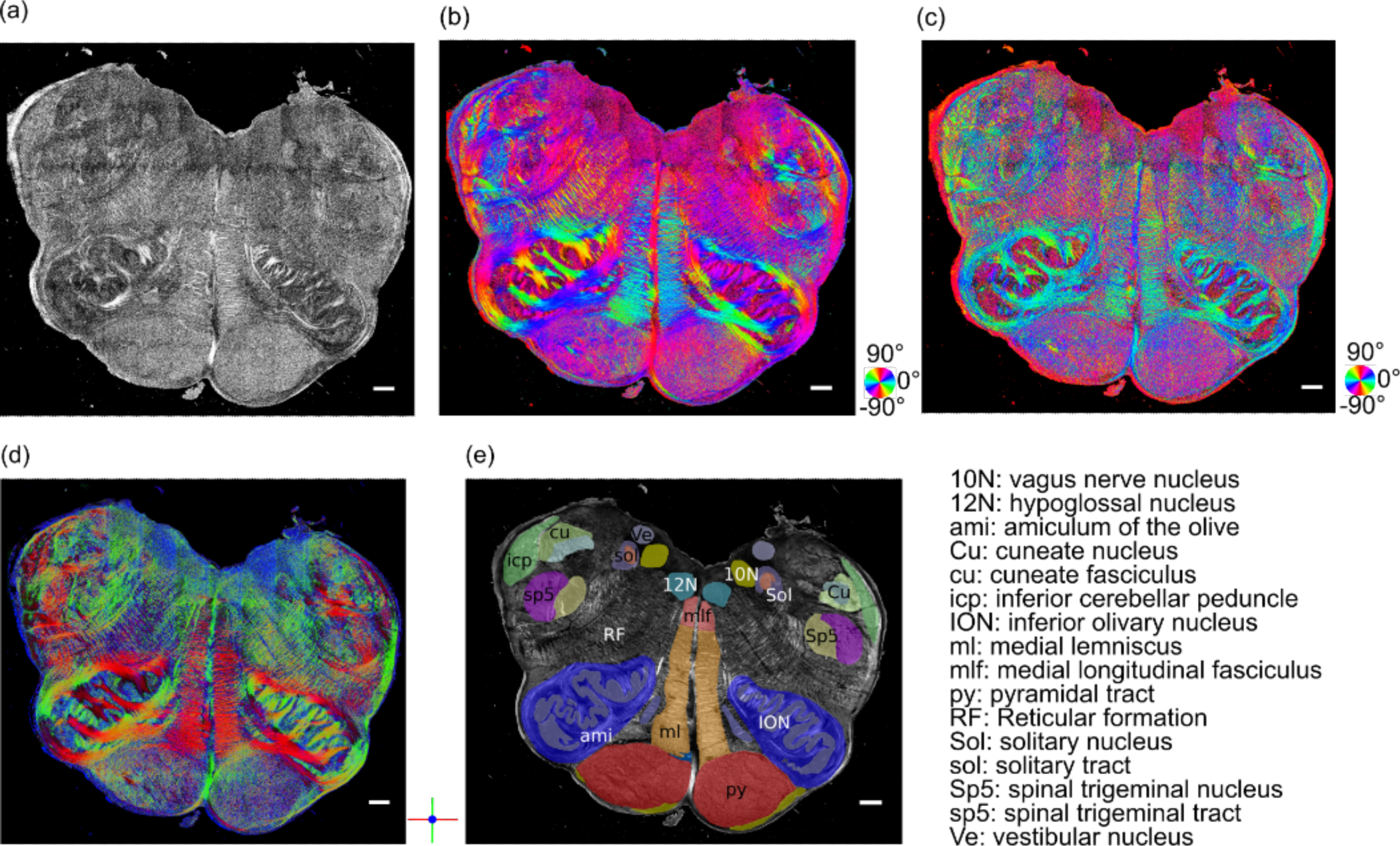
3D fiber orientation maps of a brainstem section. (a) True birefringence, (b) in-plane orientation, (c) through-plane orientation, (d) color rendering of the 3D orientation, and (e) segmentation based on the Paxinos atlas. The in-plane and through-plane orientations are color coded as indicated by the color wheels in (b) and (c). 3D orientations are color-coded as indicated by the red (left/right), green (anterior/posterior) and blue (inferior/superior) orthogonal axes in (d). Scale bars: 1mm.

We further imaged this brainstem sample in 13 consecutive block-face sections (150 µm thick each) to show the volumetric imaging capability of our method. The 1.95 mm volumetric rending of true birefringence (Supplementary Video 1), in-plane orientation (Supplementary Video 2), and through-plane orientation Supplementary Video 3) present the sophisticated fiber network in the human brainstem. Note that the dorsal region in the left medulla was missing due to imperfect tissue blocking. The microscopic details revealed by our technique can provide unprecedent information to understand the complicated fiber connections in the brainstem, for example, the connections between vagal complex with rostral arousal nuclei and diencephalon that modulate wakefulness in human consciousness ^29^.

## 4. Discussion

In this work, we presented an approach for estimating 3D fiber orientation and true birefringence of human brain samples with PS-OCT. This method formulated a birefringence vector composed of true birefringence value and 3D optic axis orientation vector, from two measurements with normal and 15 deg tilted illumination angles. We verified that our method could recover accurate estimates of a large range of through-plane angles at different in-plane orientations. We showed the 3D fiber orientation maps in full sections of a human cerebral hemisphere and a brainstem. The 10 μm resolution maps revealed intricate fiber organization beyond the capabilities of non-invasive imaging techniques such as dMRI. To our knowledge, this is the first characterization of 3D fiber axis in the human brain at microscopic spatial resolution and high angular precision, using PS-OCT based methods.

Our major contribution is the accurate and reliable estimation of 3D axis orientation in full spherical space, using only two illumination beams. Previously, quantifying 3D orientation by PS-OCT via multiple beams has been proposed based on either apparent birefringence deviation or geometrical transformation of in-plane orientations ^16,17^. A common caveat of those methods is they did not result in coverage of the full angular space, which is critical for revealing fiber configurations and structural connectivity in the human brain. Earlier studies suffered the limitation that only the axis lying within the 2D plane spanned by the illumination beams could be recovered ^17^. Later methods removed this limitation, but required a third illumination to complete the estimation in the 3D space, substantially increasing the acquisition time and hardware requirements ^16,18^. Another work utilized a dual-angle PS-OCT with a large tilt angle, based on a geometric reasoning but failed in our hands for axes co-planar to the two illumination beams ^19^. Here we significantly advanced the 3D axis orientation reconstruction with a small tilt angle of 15 deg as the second illumination, which preserves a better signal-to-noise-ratio (SNR) and simplifies image registration to the perpendicular illumination. The birefringence vector as defined in this work makes use of both apparent birefringence and in-plane orientation information in PS-OCT. The optimization formation obviates the requirement for the third illumination angle and allows the retrieval of the vector from only two sets of measurements. This implementation significantly reduces the engineering complexity and the acquisition time for large-scale samples. Importantly, we showed that our method was able to recover the axis orientation in full spherical space (Fig. 2-4) with high angular precision (Fig. 3). Therefore, the proposed PS-OCT method enables quantitative characterization of complex fiber configurations in the human brain.

Signal-to-noise-ratio (SNR) plays an important role in 3D orientation estimation. We applied refractive index matching to our samples to increase the SNR in the PS-OCT measurements. Previously, we showed that index matching significantly improved the SNR and subsequentially enhanced the accuracy of in-plane orientation and apparent birefringence measurements ^21^. It is important to note that, without index matching, the limited SNR in the data would prevent the successful application of our orientation estimation from the two illumination measurements.

Unveiling the connectional anatomy at the microscopic level is imperative for understanding the human brain. Conventional microscopy approaches involve registration of a stack of 2D histological brain sections to recover volumetric structure, a challenging task due to the nonlinear distortions induced by sectioning and tissue mounting ^30^. This renders continuous tracing of 3D fibers across sectioning at microscopic scale impractical ^3^. In this work, we showed examples of 3D axis orientation maps in full sections from human cerebrum and brainstem (Fig. 5-6), revealing their complex organization at microscopic resolution and without the aforementioned distortions. Our technique paves the way towards 3D microscopic tractography. We plan to achieve this by incorporating a serial sectioning PS-OCT technique ^13^, whose unique advantage is that the blockface of the tissue is imaged before cutting, therefore eliminating tissue distortion and preserving volumetric inter-slice alignment with microscopic accuracy ^30^. The further development of serial PS-OCT opens the potential of large-scale 3D fiber axis imaging and 3D microscopic tractography.

Diffusion MRI has been widely applied to reconstruct large fiber pathways in the human brain. However, the millimeter resolution of dMRI is insufficient in regions where multiple microscopic fascicles converge ^26^; therefore, microscopy is need to validate and to guide dMRI tractography ^3^. PS-OCT has been used to evaluate the accuracy of dMRI in-plane fiber orientation obtained with different q-space sampling schemes, spatial resolutions, and orientation reconstruction methods in human white matter samples ^31,32^. The ability to measure 3D orientation in large-scale samples will enable full validation of dMRI orientation estimates.

In this work, we also introduced true birefringence as a quantitative measure of myelin content (Fig. 2e). The alteration of myelination has been observed in many neurodegenerative diseases, including cerebral amyloid angiopathy ^33^, Alzheimer’s disease ^34^, and multiple sclerosis ^35^. The apparent birefringence obtained from conventional PS-OCT depends on the through-plane angle of the fibers (Fig. 2e), which biases the assessment of myelin. To obtain an accurate metric of myelin content, the effect of the through-plane fiber axis needs to be removed. Our method of estimating true birefringence in the brain is a potential solution to this problem that merits further investigation and validation. The ability to quantify the myelin content in tens to hundreds of cubic centimeters of brain tissue promises to advance our understanding of neuropathology in various neurodegenerative diseases.

We presented intricate 3D fiber organizations in the human brain, which are beyond the ability of current imaging tools (Fig. 5-Fig. 6). The current application on brain mapping addresses one of the most sophisticated fibrous networks, such as shown in the brainstem where small fiber tracts form a complex 3D network ^36^. It is worthwhile emphasizing that our method can be applied to quantitatively image the 3D axis orientation and true birefringence in other fibrous tissues as well, such as cartilage ^18^, capsular ligament ^37^, enamel ^38^, uterus ^39^, and heart ^19^. Characterizing the 3D fiber organization in these tissues will play an essential role to understand the normal functions and the pathological impact in diseases.

One limitation of this study is that we assumed that the axis of fibers bundles was homogenous within a small volume at the intersection of the two beams, i.e., approximately within 150 μm × sinΩ in this study. Therefore, we only applied our method on the *en-face* orientation data. In future work, we will develop methods to extract local birefringence and orientation ^19,40–42^ to further obtain depth-resolved 3D fiber orientation. We also plan to develop an automatic serial PS-OCT system with two beam illuminations for large-scale *ex vivo* human brain imaging. Combination of quantitative 3D orientation and serial sectioning PS-OCT opens new avenues towards imaging connectional anatomy in the healthy and diseased brain at unprecedented scales.

## Acknowledgement

We would like to thank Dr. Christophe Destrieux from Université de Tours body donation program for providing the brainstem sample, and Dr. Divya Varadarajan at the Martinos Center for the valuable discussions.

## Data availability

All data included in this study are presented in the figures. Raw image data of the human brain may be obtained from the corresponding author following MGH guideline.

## References

1 Van Essen, D. C. et al. The WU-Minn Human Connectome Project: An overview. Neuroimage 80, 62–79, 10.1016/j.neuroimage.2013.05.041 (2013).

2 Huang, S. Y. et al. Connectome 2.0: Developing the next-generation ultra-high gradient strength human MRI scanner for bridging studies of the micro-, meso-and macro-connectome. Neuroimage 243, 118530, 10.1016/j.neuroimage.2021.118530 (2021).

3 Yendiki, A. et al. Post mortem mapping of connectional anatomy for the validation of diffusion MRI. Neuroimage 256, 119146, 10.1016/j.neuroimage.2022.119146 (2022).

4 Wu, C. et al. A Novel Fluorescent Probe That Is Brain Permeable and Selectively Binds to Myelin. Journal of Histochemistry & Cytochemistry 54, 997–1004, doi:10.1369/jhc.5A6901.2006 (2006).

5 Farrar, Matthew J., Wise, Frank W., Fetcho, Joseph R. & Schaffer, Chris B. In Vivo Imaging of Myelin in the Vertebrate Central Nervous System Using Third Harmonic Generation Microscopy. Biophysical Journal 100, 1362–1371, doi:10.1016/j.bpj.2011.01.031 (2011).

6 Wang, H., Fu, Y., Zickmund, P., Shi, R. & Cheng, J.-X. Coherent Anti-Stokes Raman Scattering Imaging of Axonal Myelin in Live Spinal Tissues. Biophysical Journal 89, 581–591, 10.1529/biophysj.105.061911 (2005).

7 Khan, A. R. et al. 3D structure tensor analysis of light microscopy data for validating diffusion MRI. Neuroimage 111, 192–203, doi:10.1016/j.neuroimage.2015.01.061 (2015).

8 Schurr, R. & Mezer Aviv, A. The glial framework reveals white matter fiber architecture in human and primate brains. Science 374, 762–767, doi:10.1126/science.abj7960 (2021).

9 Leuze, C. et al. Comparison of diffusion MRI and CLARITY fiber orientation estimates in both gray and white matter regions of human and primate brain. Neuroimage 228, 117692–117692, doi:10.1016/j.neuroimage.2020.117692 (2021).

10 Wang, H., Lenglet, C. & Akkin, T. Structure tensor analysis of serial optical coherence scanner images for mapping fiber orientations and tractography in the brain. Journal of Biomedical Optics 20, 1–11, doi:10.1117/1.JBO.20.3.036003 (2015).

11 Menzel, M. et al. A Jones matrix formalism for simulating three-dimensional polarized light imaging of brain tissue. Journal of The Royal Society Interface 12, 20150734, doi:doi:10.1098/rsif.2015.0734 (2015).

12 Axer, M. et al. A novel approach to the human connectome: Ultra-high resolution mapping of fiber tracts in the brain. Neuroimage 54, 1091–1101, 10.1016/j.neuroimage.2010.08.075 (2011).

13 Wang, H. et al. as-PSOCT: Volumetric microscopic imaging of human brain architecture and connectivity. Neuroimage 165, 56–68, 10.1016/j.neuroimage.2017.10.012 (2018).

14 Liu, C. J. et al. Quantification of volumetric morphometry and optical property in the cortex of human cerebellum at micrometer resolution. Neuroimage 244, 118627, 10.1016/j.neuroimage.2021.118627 (2021).

15 Schmitz, D. et al. Derivation of Fiber Orientations From Oblique Views Through Human Brain Sections in 3D-Polarized Light Imaging. Frontiers in Neuroanatomy 12 (2018).

16 Liu, C. J., Black, A., Wang, H. & Akkin, T. Quantifying three-dimensional optic axis using polarization-sensitive optical coherence tomography. Journal of Biomedical Optics 21, 070501 (2016).

17 Ugryumova, N., Gangnus, S. V. & Matcher, S. J. Three-dimensional optic axis determination using variable-incidence-angle polarization-optical coherence tomography. Opt. Lett. 31, 2305–2307, doi:10.1364/OL.31.002305 (2006).

18 Ugryumova, N., Jacobs, J., Bonesi, M. & Matcher, S. J. Novel optical imaging technique to determine the 3-D orientation of collagen fibers in cartilage: variable-incidence angle polarization-sensitive optical coherence tomography. Osteoarthritis and cartilage 17, 33–42 (2009).

19 Wang, Y., Ravanfar, M., Zhang, K., Duan, D. & Yao, G. Mapping 3D fiber orientation in tissue using dual-angle optical polarization tractography. Biomed. Opt. Express 7, 3855–3870, doi:10.1364/BOE.7.003855 (2016).

20 Larsen, L., Griffin, L. D., Gräßel, D., Witte, O. W. & Axer, H. Polarized light imaging of white matter architecture. Microscopy research and technique 70, 851–863, doi:10.1002/jemt.20488 (2007).

21 Liu, C. J. et al. Refractive-index matching enhanced polarization sensitive optical coherence tomography quantification in human brain tissue. Biomed. Opt. Express 13, 358–372, doi:10.1364/BOE.443066 (2022).

22 Thirion, J. P. Image matching as a diffusion process: an analogy with Maxwell’s demons. Medical Image Analysis 2, 243–260, 10.1016/S1361-8415(98)80022-4 (1998).

23 Kroon, D. & Slump, C. H. in 2009 IEEE International Symposium on Biomedical Imaging: From Nano to Macro. 963–966.

24 Veraart, J. et al. Denoising of diffusion MRI using random matrix theory. Neuroimage 142, 394–406, 10.1016/j.neuroimage.2016.08.016 (2016).

25 Andersson, J. L. R. & Sotiropoulos, S. N. An integrated approach to correction for off-resonance effects and subject movement in diffusion MR imaging. Neuroimage 125, 1063–1078, 10.1016/j.neuroimage.2015.10.019 (2016).

26 Mori, S., Crain, B. J., Chacko, V. P. & Van Zijl, P. C. M. Three-dimensional tracking of axonal projections in the brain by magnetic resonance imaging. Annals of Neurology 45, 265–269, 10.1002/1531-8249(199902)45:2<265::AID-ANA21>3.0.CO;2-3 (1999).

27 Schmahmann, J. D. & Pandya, D. N. Fiber Pathways of the Brain. (Oxford University Press, 2006).

28 Mai, J. K. & Paxinos, G. The human nervous system. (Academic press, 2011).

29 Edlow, B. L., McNab, J. A., Witzel, T. & Kinney, H. C. The Structural Connectome of the Human Central Homeostatic Network. Brain Connectivity 6, 187–200, doi:10.1089/brain.2015.0378 (2015).

30 Ali, S. et al. Rigid and non-rigid registration of polarized light imaging data for 3D reconstruction of the temporal lobe of the human brain at micrometer resolution. Neuroimage 181, 235–251, 10.1016/j.neuroimage.2018.06.084 (2018).

31 Jones, R. et al. Insight into the fundamental trade-offs of diffusion MRI from polarization-sensitive optical coherence tomography in ex vivo human brain. Neuroimage 214, 116704, 10.1016/j.neuroimage.2020.116704 (2020).

32 Jones, R. et al. High-fidelity approximation of grid-and shell-based sampling schemes from undersampled DSI using compressed sensing: Post mortem validation. Neuroimage 244, 118621, 10.1016/j.neuroimage.2021.118621 (2021).

33 Salat, D. H. et al. White Matter Alterations in Cerebral Amyloid Angiopathy Measured by Diffusion Tensor Imaging. Stroke 37, 1759–1764, doi:10.1161/01.STR.0000227328.86353.a7 (2006).

34 Nasrabady, S. E., Rizvi, B., Goldman, J. E. & Brickman, A. M. White matter changes in Alzheimer’s disease: a focus on myelin and oligodendrocytes. Acta Neuropathologica Communications 6, 22, doi:10.1186/s40478-018-0515-3 (2018).

35. Lubetzki, C. & Stankoff, B. in Handbook of Clinical Neurology Vol. 122 (ed Douglas S. Goodin) 89-99 (Elsevier, 2014).

36 Sporns, O., Tononi, G. & Kötter, R. The Human Connectome: A Structural Description of the Human Brain. PLOS Computational Biology 1, e42, doi:10.1371/journal.pcbi.0010042 (2005).

37 Zarei, V., Liu, C. J., Claeson, A. A., Akkin, T. & Barocas, V. H. Image-based multiscale mechanical modeling shows the importance of structural heterogeneity in the human lumbar facet capsular ligament. Biomechanics and Modeling in Mechanobiology 16, 1425–1438, doi:10.1007/s10237-017-0896-4 (2017).

38 Tang, P. et al. Local axis orientation mapped by polarization sensitive optical coherence tomography provides a unique contrast to identify caries lesions in enamel. Biomed. Opt. Express 13, 4247–4260, doi:10.1364/BOE.464707 (2022).

39 Li, W., Narice, B. F., Anumba, D. O. & Matcher, S. J. Polarization-sensitive optical coherence tomography with a conical beam scan for the investigation of birefringence and collagen alignment in the human cervix. Biomed. Opt. Express 10, 4190–4206, doi:10.1364/BOE.10.004190 (2019).

40 Fan, C. & Yao, G. Mapping local optical axis in birefringent samples using polarization-sensitive optical coherence tomography. Journal of Biomedical Optics 17, 1–3, doi:10.1117/1.JBO.17.11.110501 (2012).

41 Li, Q. et al. Robust reconstruction of local optic axis orientation with fiber-based polarization-sensitive optical coherence tomography. Biomed. Opt. Express 9, 5437–5455, doi:10.1364/BOE.9.005437 (2018).

42 Tang, P. et al. Polarization sensitive optical coherence tomography with single input for imaging depth-resolved collagen organizations. Light: Science & Applications 10, 237, doi:10.1038/s41377-021-00679-3 (2021).

