## Supplemental materials for "Quantitative imaging of three-dimensional fiber orientation in the human brain via two illumination angles using polarization-sensitive optical coherence tomography"

Supplementary material

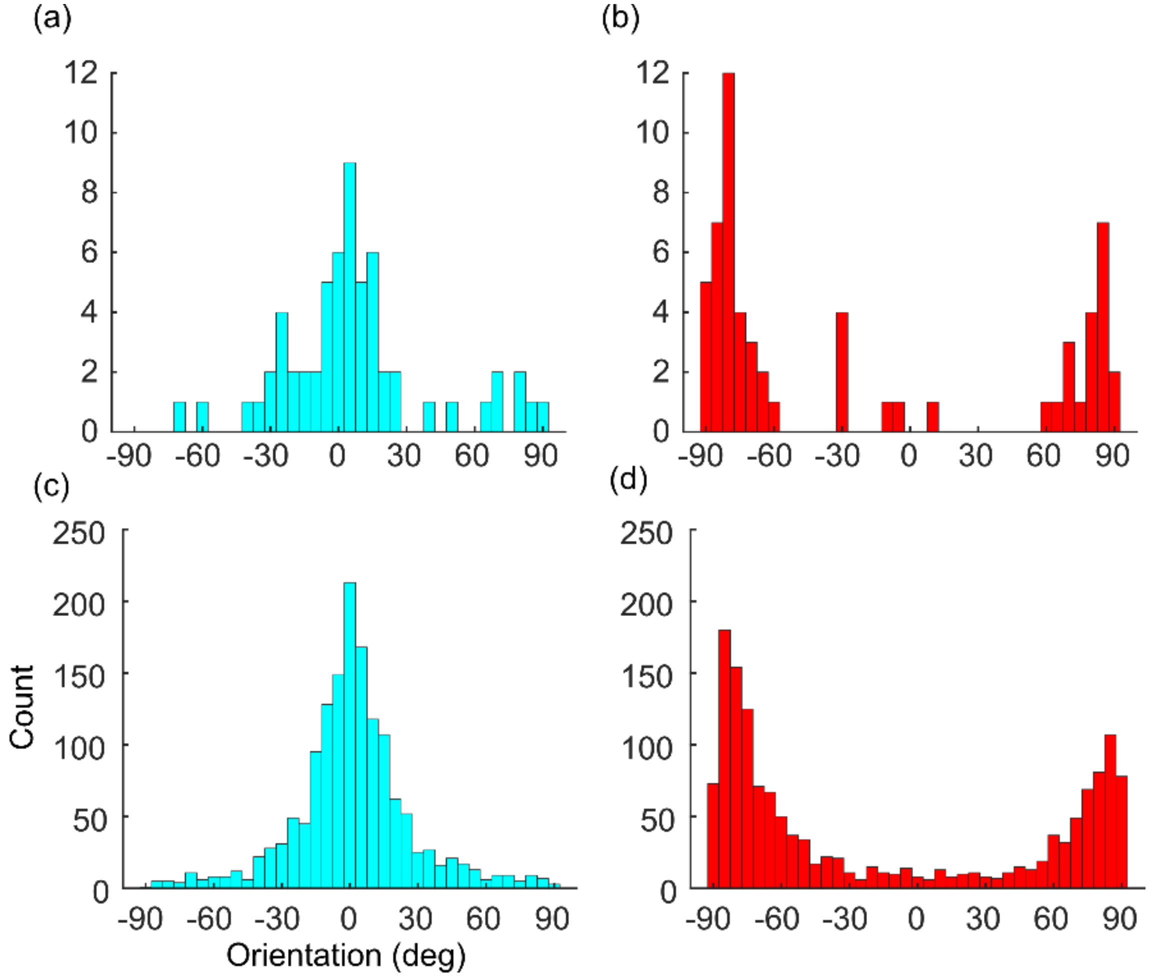


Fig. S1 Rationale of the interchange of the axis

The histogram of optic axis orientation data along one A-line (60 pixels) whose dominate orientation is 0 deg (a) and 90 deg (b). For fibers whose in-plane orientation is close to 0 deg, the input for the optimization is not affected by the noise distribution in the data. However, the U-shape distribution of optic axis orientation at 90 deg causes the sub-optimal result of the optimization without interchanging the xy axis. By binning the neighboring 25 A-lines together (1500 pixels in total), this distribution is more obvious for axis orientation at 0 deg (c) and 90 deg (d). The binning width in (a)-(d) is 5 deg.

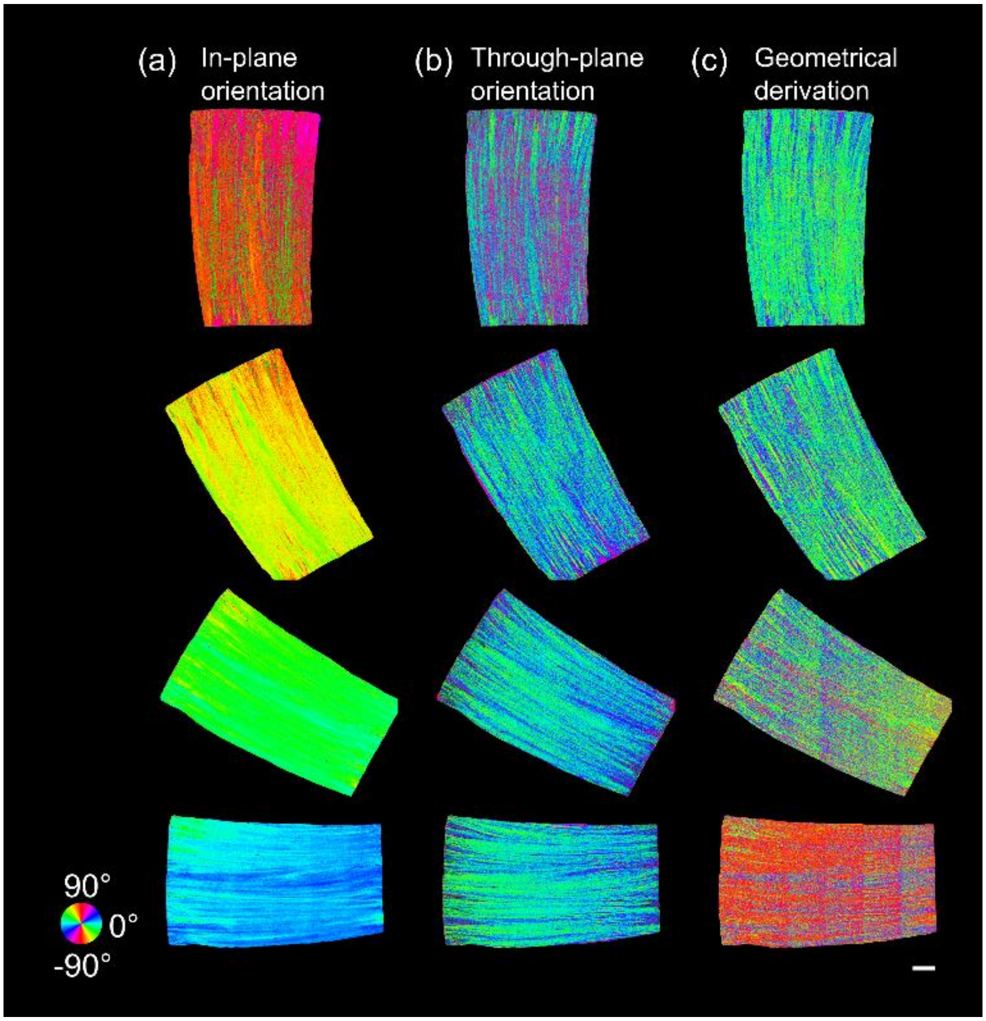


Fig. S2 Validation of the xy-axis swapping strategy in Section 2.1.
The in-plane (a) and through-plane (b) orientation images of the sample in Fig. 4 without swapping the xy axis (using Eq. 1, 2, 3 for all four in-plane rotation angles from -90 deg to 0 deg). The in-plane and through-plane orientation shows considerable errors when the degree of rotation is -90 deg (green and magenta-purple pixels in the first row of (a) and (b), respectively). (c) The through-plane orientation images derived from geometrical definitions as discussed in detail in our previous work (see Ref. 16). The equation used here is $\Psi = arccot(\frac{sin\Theta_1}{sin\Omega \tan\Theta_3} - \frac{cos\Theta_1}{tan\Omega})$, where $\Psi$ is the through-plane orientation, $\Theta_1$ is the orientation measurement from the normal illumination, $\Theta_3$ is the orientation measurement from the tilted illumination about y-axis, and $\Omega$ is the tilted angle (15 deg). The calculation is unreliable when $\Theta_1$ is close to 0 deg. The in-plane and through-plane orientations are color-coded as indicated by the color wheels. Scale bar: 1mm.
