## Supplementary figures and images for "Quantitative imaging of three-dimensional fiber orientation in the human brain via two illumination angles using polarization-sensitive optical coherence tomography"

### Supplemental video 1

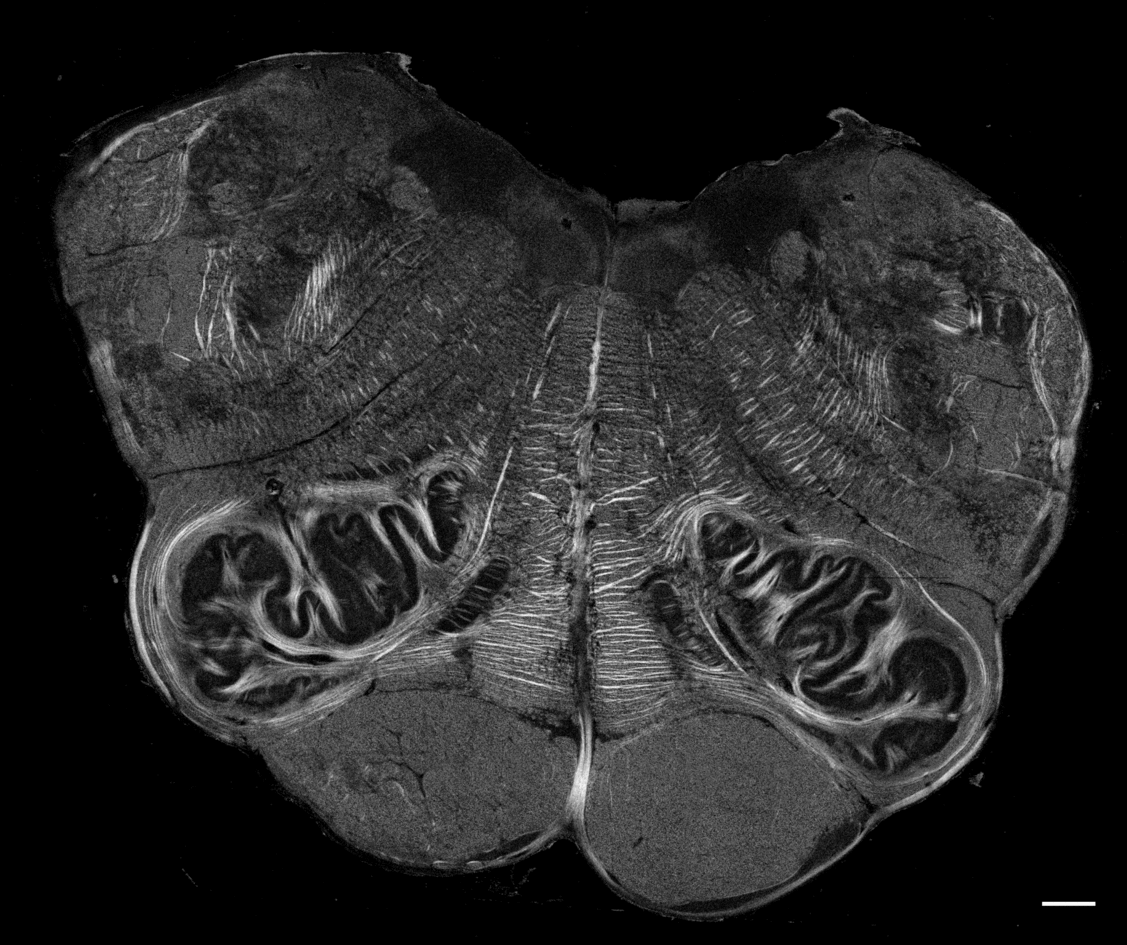

### Supplemental video 2

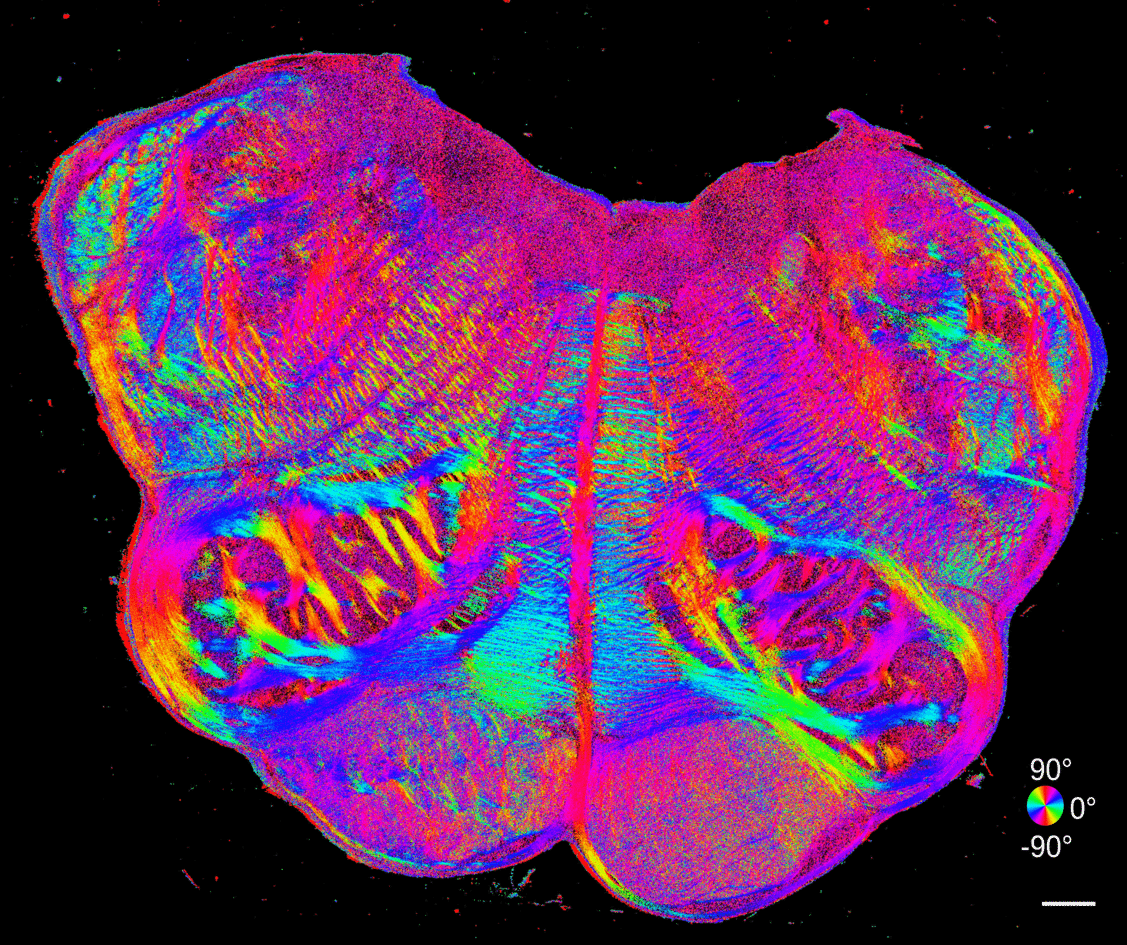

### Supplemental video 3

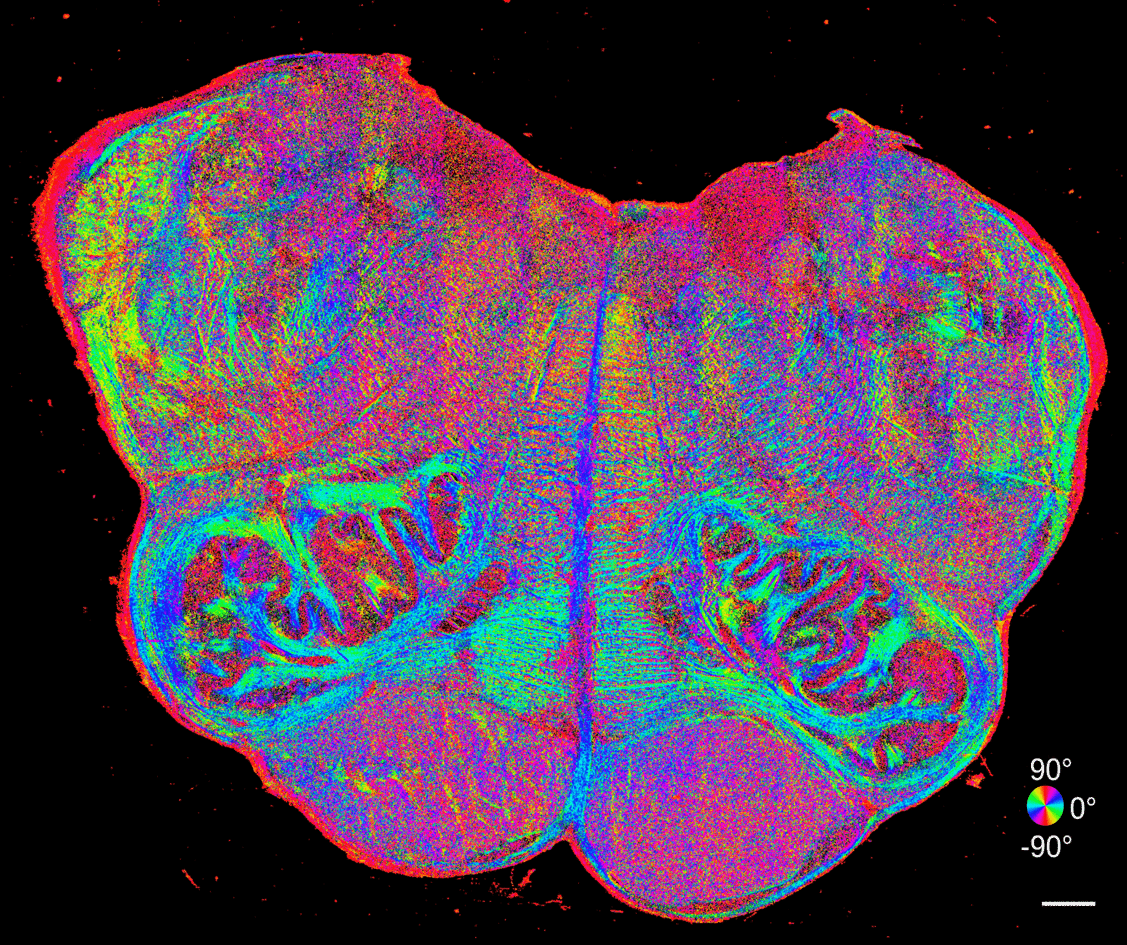
